# The histone methyltransferase DOT1L prevents antigen-independent differentiation and safeguards epigenetic identity of CD8^+^ T cells

**DOI:** 10.1101/826255

**Authors:** Eliza Mari Kwesi-Maliepaard, Muhammad Assad Aslam, Mir Farshid Alemdehy, Teun van den Brand, Chelsea McLean, Hanneke Vlaming, Tibor van Welsem, Tessy Korthout, Cesare Lancini, Sjoerd Hendriks, Tomasz Ahrends, Dieke van Dinther, Joke M.M. den Haan, Jannie Borst, Elzo de Wit, Fred van Leeuwen, Heinz Jacobs

**Author notes:** These authors contributed equally to this work (Eliza Mari Kwesi-Maliepaard, Muhammad Assad Aslam, Mir Farshid Alemdehy). Equal contribution and corresponding authors. Lead contact: Fred van Leeuwen.

## Abstract

Cytotoxic T-cell differentiation is guided by epigenome adaptations but how epigenetic mechanisms control lymphocyte development has not been well defined. Here we show that the histone methyltransferase DOT1L, which marks the nucleosome core on active genes, safeguards normal differentiation of CD8^+^ T cells. T-cell specific ablation of *Dot1L* resulted in loss of naïve CD8^+^ T cells and premature differentiation towards a memory-like state, independent of antigen exposure and in a cell-intrinsic manner. Without DOT1L, the memory-like CD8^+^ cells fail to acquire full effector functions *in vitro* and *in vivo*. Mechanistically, DOT1L controlled T-cell differentiation and function by ensuring normal T-cell receptor density and signaling, and by maintaining epigenetic identity, in part by indirectly supporting the repression of developmentally-regulated genes. Through our study DOT1L is emerging as a central player in physiology of CD8^+^ T cells, acting as a barrier to prevent premature differentiation and supporting the licensing of the full effector potential of cytotoxic T cells.

## Introduction

Lymphocyte development and differentiation is tightly regulated and provides the basis for a functional adaptive immune system. Development of mature T cells initiates in the thymus with progenitor T cells that have to pass two key checkpoints: T cell receptor (TCR) β selection and positive selection, both of which are controlled by intricate signaling pathways involving the pre-TCR/CD3 and αβTCR/CD3 complexes, respectively ^1^. Upon positive selection, mature thymocytes are licensed to emigrate and populate peripheral lymphatic organs as naïve T cells (T_N_). Further differentiation of naïve T cells into effector or memory T cells normally depends on TCR-mediated antigen recognition and stimulation. However, it has become evident that a substantial fraction of mature CD8^+^ T cells acquires memory-like features independent of exposure to foreign antigen. The origin and functionality of these unconventional memory cells in mice and humans, also referred to as innate or virtual memory cells, are only just being uncovered ^2–4^.

The dynamic changes during development and differentiation of CD8^+^ T cells are governed by transcriptional and epigenetic changes, including histone modifications that are controlled by chromatin modifiers. Well-established histone marks are mono- and tri-methylation of histone H3K4 at enhancers (H3K4me1) and promoters (H3K4me3), H3K27me3 at repressed promoters, and H3K9me2/3 in heterochromatin ^5–10^. Although epigenetic ‘programming’ is known to play a key role in T cell development and differentiation, the causal role of epigenetic modulators in T cell differentiation is still poorly understood, especially for chromatin modifiers associated with active chromatin ^5^.

One of the histone modifications positively associated with gene activity is mono-, di- and tri-methylation of histone H3K79 mediated by DOT1L. This evolutionarily conserved histone methyltransferase methylates H3K79 in transcribed promoter-proximal regions of active genes ^11, 12^. Although the association with gene activity is strong, how H3K79 methylation affects transcription is still unclear and repressive functions have also been proposed ^13–18^. DOT1L has been linked to several critical cellular functions, including embryonic development, DNA damage response, and meiotic checkpoint control ^19, 20^ and DOT1L has also been shown to function as a barrier for cellular reprogramming in generating induced pluripotent stem cells ^21^. DOT1L gained wide attention as a specific drug target in the treatment of MLL-rearranged leukemia, where MLL fusion proteins aberrantly recruit DOT1L to MLL target genes leading to their enhanced expression ^22–24^. A similar dependency on DOT1L activity and sensitivity to DOT1L inhibitors was recently observed in thymic lymphoma ^25^. Interestingly, inhibition of DOT1L activity in human T cells attenuates graft-versus-host disease in adoptive cell transfer models ^26^ and it regulates CD4^+^ T cell differentiation ^27^.

Given the emerging role of DOT1L in epigenetic reprogramming and T-cell malignancies, we investigated the role of DOT1L in normal T cell physiology using a mouse model in which *Dot1L* was selectively deleted in the T cell lineage. Our results suggest a model in which DOT1L plays a central role in CD8^+^ T cell differentiation, acting as a barrier to prevent premature antigen-independent differentiation, and promoting the licensing of the full effector potential by maintaining epigenetic integrity.

## Results

### DOT1L prohibits premature differentiation of naïve towards memory-like CD8^+^ T cells

Given the essential role of DOT1L in embryonic development ^28^, we determined the role of DOT1L in T cell development and differentiation by employing a conditional knock-out mouse model in which *Dot1L* is deleted in the T-cell lineage by combining floxed *Dot1L* with a Cre-recombinase under the control of the Lck promoter, which leads to deletion of exon 2 of *Dot1L* during early thymocyte development (Sup Fig. 1a) ^25^. The observed global loss of H3K79me2 in *Lck-*Cre*^+/−^;Dot1L^fl/fl^* mice, as confirmed by immunohistochemistry on fixed thymus tissue (Sup Fig. 1b), agreed with the notion that DOT1L is the sole methyltransferase for H3K79 ^11, 25, 28, 29^.

To validate the efficacy of *Dot1L* deletion at the single-cell level, we developed an intracellular staining protocol for H3K79me2. Histone dilution by replication-dependent and -independent means has been suggested to be the main mechanisms of losing methylated H3K79 ^15, 30–32^. Flow-cytometric analyses of thymocyte subsets from *Lck-*Cre*^+/−^;Dot1L^fl/fl^* mice (hereafter, KO) revealed that double-negative (DN, CD4^−^ CD8^−^) thymocytes started losing H3K79me2. From the subsequent immature single positive state (ISP) onward all the thymocytes had lost DOT1L mediated H3K79me (Sup Fig. 1c). This confirmed that upon early deletion of *Dot1L*, successive rounds of replication in the thymus allowed for loss of methylated H3K79.

No changes in H3K79 methylation levels were found in T-lineage cells of *Lck-*Cre*^+/−^;Dot1L^wt/wt^* control mice (hereafter, WT). Ablation of *Dot1L* did not significantly affect the overall thymic cellularity, however it led to a reduction in the number of mature single positive (SP) CD4^+^ and CD8^+^ thymocytes (both 2.5-fold) while the ISP and CD4^+^CD8^+^ double positive (DP) subsets were not significantly affected, suggesting a role of DOT1L in controlling intrathymic T cell selection (Sup Fig. 1d-e).

In the spleen, overall cellularity was also not affected but within the T cell compartment, CD4^+^ T cells were drastically reduced (3.2-fold), whereas CD8^+^ T cells were increased (1.7-fold) in number (Sup Fig. 1e-f). However, while flow cytometry of H3K79me2-stained splenic T cells confirmed the lack of DOT1L activity in CD8^+^ T cells and CD44^−^CD62L^+^ CD4^+^ T cells, CD44^+^CD62L^−^ CD4^+^ cells showed a partial loss of H3K79me2 and CD4^+^CD25^+^ regulatory T cells (Treg) remained H3K79me2 positive (Sup Fig. 1g). Since earlier in development, CD4-expressing cells in the thymus were mostly H3K79me2 negative, this suggests that a strong selection occurred for the maintenance of DOT1L for the development of Tregs in this mouse model. Indeed, partial deletion of *Dot1L* in CD4^+^ cells was confirmed by PCR analysis (Sup Fig. 1h). Here, we focused our study on defining the role of DOT1L in the cytotoxic T cell compartment in which efficient deletion of *Dot1L* and loss of H3K79me2 was found in both the thymus and the periphery.

CD8^+^ T cell differentiation was strongly affected by the absence of DOT1L. Analysis of CD8^+^ T cell subsets in the spleen revealed that *Dot1L*-KO mice showed a severe loss of naïve (CD44^−^CD62L^+^) CD8^+^ T (T_N_) cells which was linked to a massive gain of the CD44^+^CD62L^+^ phenotype, a feature of central memory T cells (T_CM_; Fig. 1A-B*). Lck-*Cre*^+/−^;Dot1L^fl/wt^* heterozygous knock-out mice (Het) did not show any phenotypic differences of CD8^+^ T cells (Fig. 1A-B). The lack of haplo-insufficiency was further confirmed by principal component analysis of RNA-Seq data indicating that WT and Het CD8^+^ T cells clustered together but were separated from the KO CD8^+^ T cells, excluding gene-dosage effects (Sup Fig. 1i). Therefore, we restricted our further studies to the comparison of the KO and WT mice. The strong shift towards a CD8^+^ memory-phenotype in *Dot1L* KO was unexpected because *Dot1L*-KO mice were housed under the same conditions as their WT controls and the mice had not been specifically immunologically challenged.

**Figure 1:**
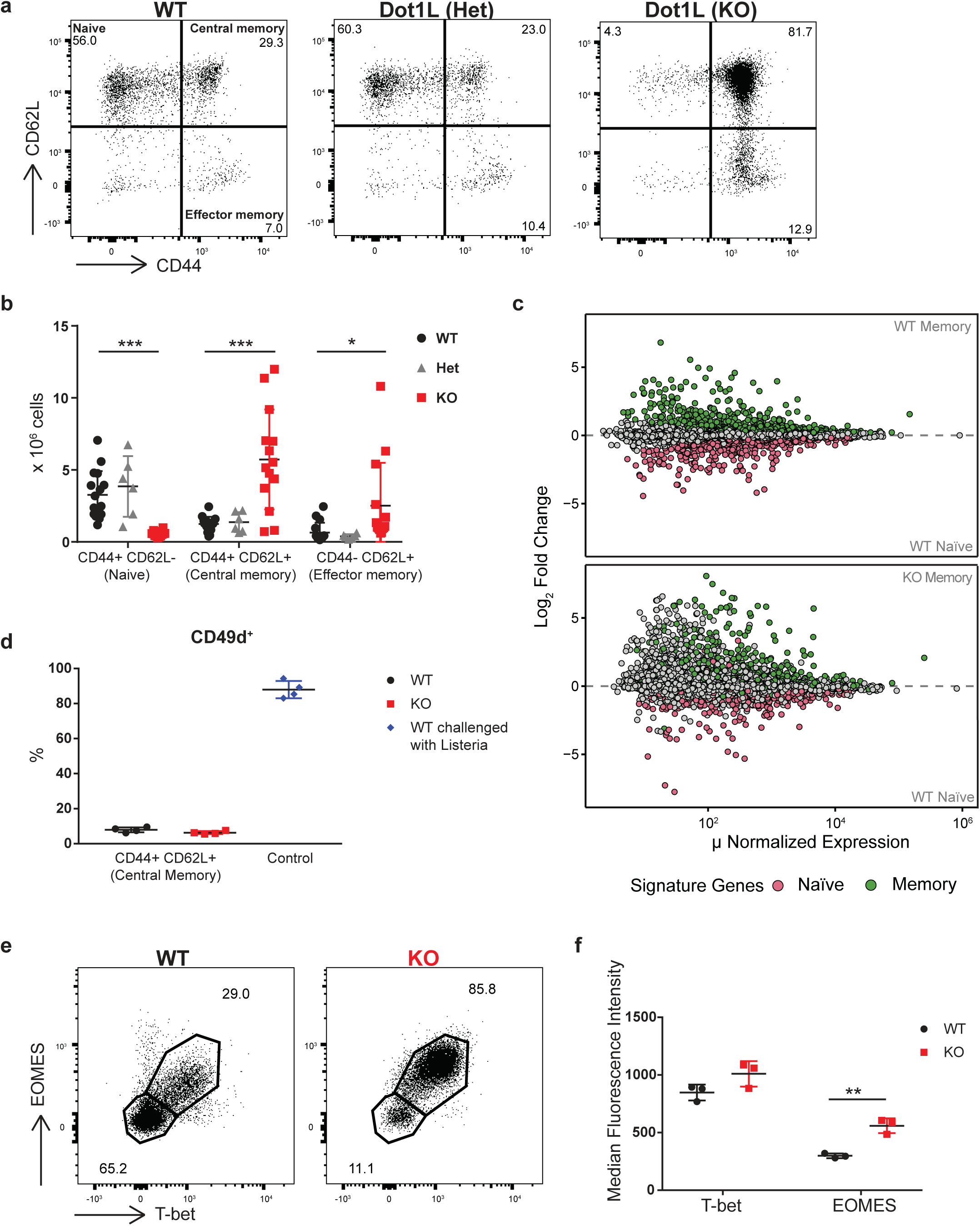
Ablation of *Dot1L* results in CD44^+^CD62L^+^ memory-like CD8^+^ T cells. a) Analysis of CD8^+^ T cell subsets in the spleen based on CD44 and CD62L expression in WT and heterozygous and homozygous *Dot1L*-KO mice. Subsets are indicated in the top left panel. b) Quantification of CD8^+^ T cell subsets in the spleen indicated in the representative plots in panel a. Data from 3-5 individual experiments with 3-4 mice per genotype per experiment, shown as mean ±SD. c) RNA-Seq analysis of sorted CD44^+^CD62L^+^ (T_CM_) and CD44^−^CD62L^+^ (T_N_) CD8^+^ T cells from four mice per indicated genotype. Naïve and memory signatures were defined based on differentially expressed genes (FDR < 0.01) between WT T_N_ and WT T_CM_ cells. d) Percentage of CD49d^+^ cells in CD44^+^CD62L^+^ CD8^+^ T cells from unchallenged mice and in CD44^+^CD62L^−^ CD8^+^ T cells from WT mice challenged with L. monocytogenes for 7 days. Data from one experiment with four mice per genotype, represented as mean ±SD. e) Representative flow-cytometry plots of T-bet and EOMES expression in CD8^+^ T cells from the spleen. f) Median fluorescence intensity of T-bet and EOMES in CD44^+^CD62L^+^ CD8^+^ T cells from the spleen. Data from one experiment with three mice per genotype, represented as mean ±SD.

To unravel the molecular identity of the CD44^+^CD62L^+^ *Dot1L*-KO cells in more detail we performed RNA-Seq analysis on sorted CD44^−^CD62L^+^ (T_N_) and CD44^+^CD62L^+^ (T_CM_) CD8^+^ T cell subsets from WT and KO mice. Based on differential gene expression between T_N_ and T_CM_ CD8^+^ cells from WT mice, naïve and memory gene signatures were defined. Interestingly, overlay of these signatures on WT CD44^−^CD62L^+^ (T_N_) and KO CD44^+^CD62L^+^ (T_CM_) cells showed that differential expression between T_N_ and T_CM_ cells was mostly conserved when *Dot1L* was ablated. (Fig. 1c), although there was misregulation of other genes as well (see below). These data suggest that in the absence of DOT1L, CD8^+^ T cells cannot retain the naïve identity but rather acquire, prematurely and in the absence of any overt immunological challenge, a transcriptome of memory-like CD8^+^ T cells.

### *Dot1L* KO memory-like cells are antigen inexperienced

Although WT and KO mice were exposed to the same environment it cannot be excluded that KO mice responded differentially to antigens in the environment. If this is the case one expects skewing in the clonality of the TCRβ gene usage. In order to investigate this possibility, we examined the TCRβ repertoire. *Tcrb*-sequencing revealed no difference in productive clonality scores between WT and KO CD8^+^ T cells (Sup Fig. 2a). Also, CDR3 length as well as *Tcrb-V* and *Tcrb-J* gene usage were unaffected (Sup Fig. 2b-d). These data, together with the nearly complete loss of naïve CD8^+^ T cells, argued against any antigen-mediated bias in the selection for CD8^+^ T cells and indicated that CD44^+^CD62L^+^ memory-like CD8^+^ T cells in KO mice were polyclonal and arose by antigen-independent differentiation of T_N_ cells.

Antigen-independently differentiated memory-like CD8^+^ T cells have already been described in the literature and their origins and functions are subject of ongoing studies ^2–4, 33^. Depending on their origin and cytokine dependency they are referred to as ‘virtual’ or ‘innate’ memory cells. Virtual memory CD8^+^ T cells have been suggested to arise in the periphery from cells that are CD5^high^, related to high TCR affinity, and require IL-15 ^4, 34^. In contrast, innate memory CD8^+^ T cells develop in the thymus and their generation and survival are generally considered to be dependent on IL-4 signaling ^3^. We here collectively refer to them as Antigen-Independent Memory-like CD8^+^ T cells (T_AIM_). A common feature of these T_AIM_ cells is reduced expression of CD49d ^35^, a marker that is normally upregulated after antigen exposure. In addition, they express high levels of T-bet and EOMES, encoding two memory/effector transcription factors ^36, 37^. In both *Dot1L-*KO and WT, the majority of the CD44^+^CD62L^+^ (T_CM_) cells were CD49d negative. As a control, CD44^+^CD62L^−^ effector T cells (T_EFF_) from WT mice challenged with *Listeria monocytogenes* were mostly CD49d positive (Fig. 1d). This further indicates that the generation of CD44^+^CD62L^+^ memory-like T cells in KO mice is independent of antigen exposure. Of note, the percentage of CD49d-negative CD44^+^CD62L^+^ cells that we observed in WT mice corresponds to the percentage of T_AIM_ cells reported in WT B6 mice ^38^. Regarding the expression of T-bet and EOMES, most of the KO CD8^+^ T cells co-expressed T-bet and EOMES. Furthermore, EOMES was expressed at a higher level in KO CD8^+^ T cells as compared to their WT counterpart (Fig. 1e-f). Together these characteristics are all in agreement with antigen-independent differentiation of naïve CD8^+^-T cells in the absence of DOT1L.

### The T_AIM_ phenotype in *Dot1L* KO initiates in the thymus and is cell-intrinsic

To determine whether peripheral T_AIM_ cells observed in the *Dot1L*-KO setting originate intrathymically, as reported previously for IL-4 dependent innate memory T cells ^3^, we compared RNA-Seq data from SP CD8^+^ thymocytes from KO and WT mice. Analyzing the relative distribution of memory and naïve signature genes revealed that memory genes were among the genes upregulated in KO SP CD8^+^ thymocytes (Fig. 2a). Importantly, like in peripheral CD8^+^ T cells the expression of *T-bet* and *Eomes* was highly upregulated in KO SP CD8^+^ thymocytes. This transcriptional upregulation was corroborated by flow cytometric analysis of protein expression. Intracellular staining for the transcription factors showed that a small but substantial subset of the SP CD8^+^ KO thymocytes expressed T-bet and EOMES at the protein level (Fig. 2b-c). Together with the unperturbed TCRβ repertoire, this further supports the notion that differentiation of *Dot1L*-KO CD8^+^ T cells towards memory-like cells initiates intrathymically in an antigen-independent manner. Innate memory cells have been suggested to arise in the thymus in response to an increase in IL-4 producing PLZF^high^ invariant NKT (iNKT) cells or γδ T cells ^3^. However, iNKT cells were nearly absent in the thymus of *Dot1L-*KO mice (Fig. 2d-e). Furthermore, the number of γδ T cells did not differ significantly between WT and KO mice (Fig. 2f). Previous studies on innate memory T cells from different mouse models demonstrated that introduction of a transgenic TCR, inhibiting the generation of iNKT and γδ T cells, prevents the development of innate memory cells ^39, 40^. However, introduction of the transgenic OT-I TCR, a condition under which the number of iNKT and γδ T cells is strongly reduced ^41, 42^, did not affect the memory-phenotype of *Dot1L*-KO CD8^+^ T cells (Fig. 2g). Together, these findings indicate that the intrathymic differentiation of T_AIM_ CD8^+^ cells in the absence of DOT1L did not depend on an excess of IL4-producing cells in the thymic microenvironment as reported for innate memory T cells. Rather the formation of T_AIM_ cells in *Dot1L-*KO mice likely relates to a cell-intrinsic mechanism.

**Figure 2:**
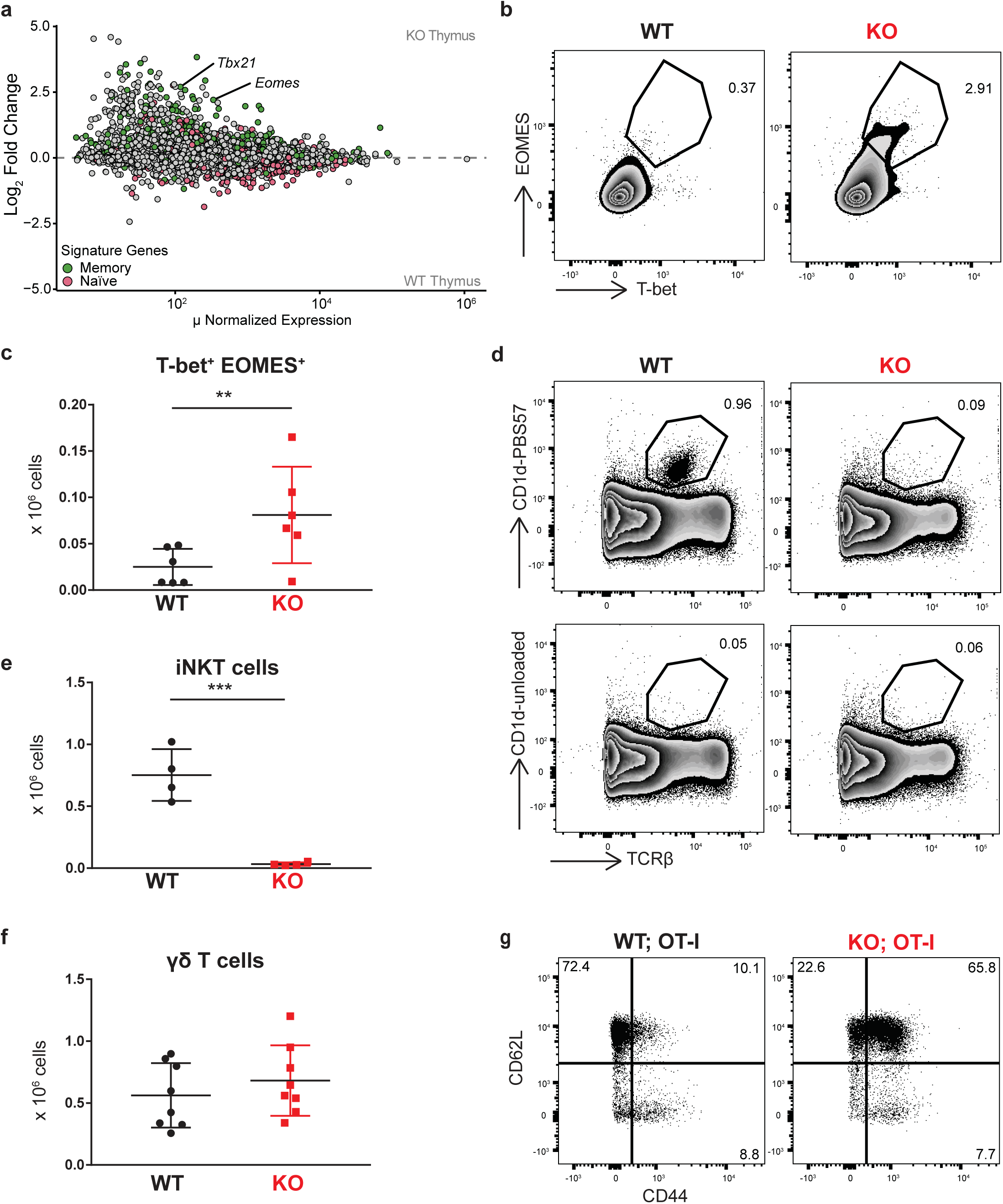
The T_AIM_ phenotype in *Dot1L* KO initiates in the thymus and is cell-intrinsic. a) MA-plot of RNA-Seq data from sorted CD4^−^CD8^+^CD3^+^ thymocytes from three WT and four KO mice. Naïve and memory signature genes were defined as described in Fig. 1C. b) Representative plot and c) quantification of T-bet and EOMES expression on CD4^−^CD8^+^CD3^+^ thymocytes, data of two individual experiments with three mice per genotype, represented as mean ±SD. d-e) Representative plots and quantification of iNKT (CD1d-PBS57^+^TCRβ^+^) cells in total thymus. Data from one experiment with four mice per genotype, represented as mean ±SD. f) Absolute number of γδTCR^+^ cells in spleen, data of two individual experiments with four mice per genotype, represented as mean ±SD. g) Representative plot of CD44 and CD62L expression on CD8^+^ T cells in *Lck-*Cre;*Dot1L;OT-I* mice.

### *Dot1L* ablation impairs TCR/CD3 expression

One of the cell-intrinsic mechanisms reported to be involved in the formation of T_AIM_ cells is aberrant TCR signaling ^2, 34, 43, 44^. Furthermore, treatment of human T cells with DOT1L inhibitor impaired TCR sensitivity and attenuated low avidity T cell responses ^26^. This led us to investigate the expression of genes encoding TCR signaling components in the absence of DOT1L. RNA-Seq analyses confirmed that many TCR signaling genes were differentially expressed between WT and KO SP CD8^+^ thymocytes (Fig. 3a). Importantly, CD3ζ (CD247), a critical rate limiting factor in controlling the transport of fully assembled TCR/CD3 complexes to the cell surface, was downregulated in *Dot1L*-KO T cells ^1, 45–48^. In addition, other components of the TCR/CD3 complex like CD3e and its associated co-receptor CD8a/b were also downregulated in KO T cells. H3K79me2 ChIP-Seq showed that these genes had H3K79me2 in WT mice and might therefore be directly regulated by DOT1L (Sup Fig. 3a-c). As a consequence, one expects TCR/CD3 and CD8αβ to be reduced at the cell surface of T_AIM_ cells, which we confirmed by flow cytometry (Fig. 3b). In addition to the CD3/TCR complex, we observed downregulation of *Itk*, a key TCR signaling molecule reported to be involved in innate memory CD8^+^ T cell formation ^49^. To exclude that the impaired TCR signaling in KO T cells could be compensated for by the selection of thymocytes expressing TCRs with altered affinity, we kept the TCR affinity identical by crossing the OT-I TCR transgene into our system. Thymocytes expressing OT-I are positively selected in the presence of MHC class I (H-2K^b^), mainly generating CD8^+^ SP T cells expressing the exogenous OT-I TCR, with concomitant reduction of the CD4^+^ lineage ^50^. If DOT1Ldeficiency impairs TCR surface density and signaling, positive selection of conventional OT-I CD8 thymocytes is expected to be compromised. In *Dot1L-*KO mice expressing OT-I, the number of SP CD8^+^ thymocytes was decreased (3.8-fold) compared to WT mice expressing OT-I (Fig. 3c). This revealed that like with endogenous TCR, early intrathymic ablation of *Dot1L* in the T cell lineage prohibits positive section of conventional OT-I CD8^+^ T cells, but yet supports the generation and selection of T_AIM_ cells (Fig. 2g) of which the vast majority expressed both exogenous TCR chains (TCRαV2 and TCRβV5) (Fig. 3d). Consistent with the low surface expression of TCR/CD3 and CD8 in the KO condition, the surface expression of OT-I TCR was also lower, as determined by SIINFEKL/H-2K^b^ tetramer staining (Sup Fig. 3d). This was further validated by staining with antibodies for the transgenic TCR chains TCRαV2 and TCRβV5 which indicated two-fold reduction in KO (Fig. 3b and d). The TCRαV2 element of the OT-I TCR was under the control of an exogenous promoter, suggesting that the reduced surface expression of OT-I TCR on KO T cells did not relate to transcriptional silencing of native TCR gene promoters. Instead, the observed lower TCR surface levels in KO T cells likely relate to the reduced CD3ζ expression ^48^. In conclusion, DOT1L regulates the levels of TCR complex and signaling molecules independent of the selected TCR. In the absence of these regulatory mechanisms, the identity of naïve CD8^+^ T cells cannot be maintained, contributing to premature T cell differentiation.

**Figure 3:**
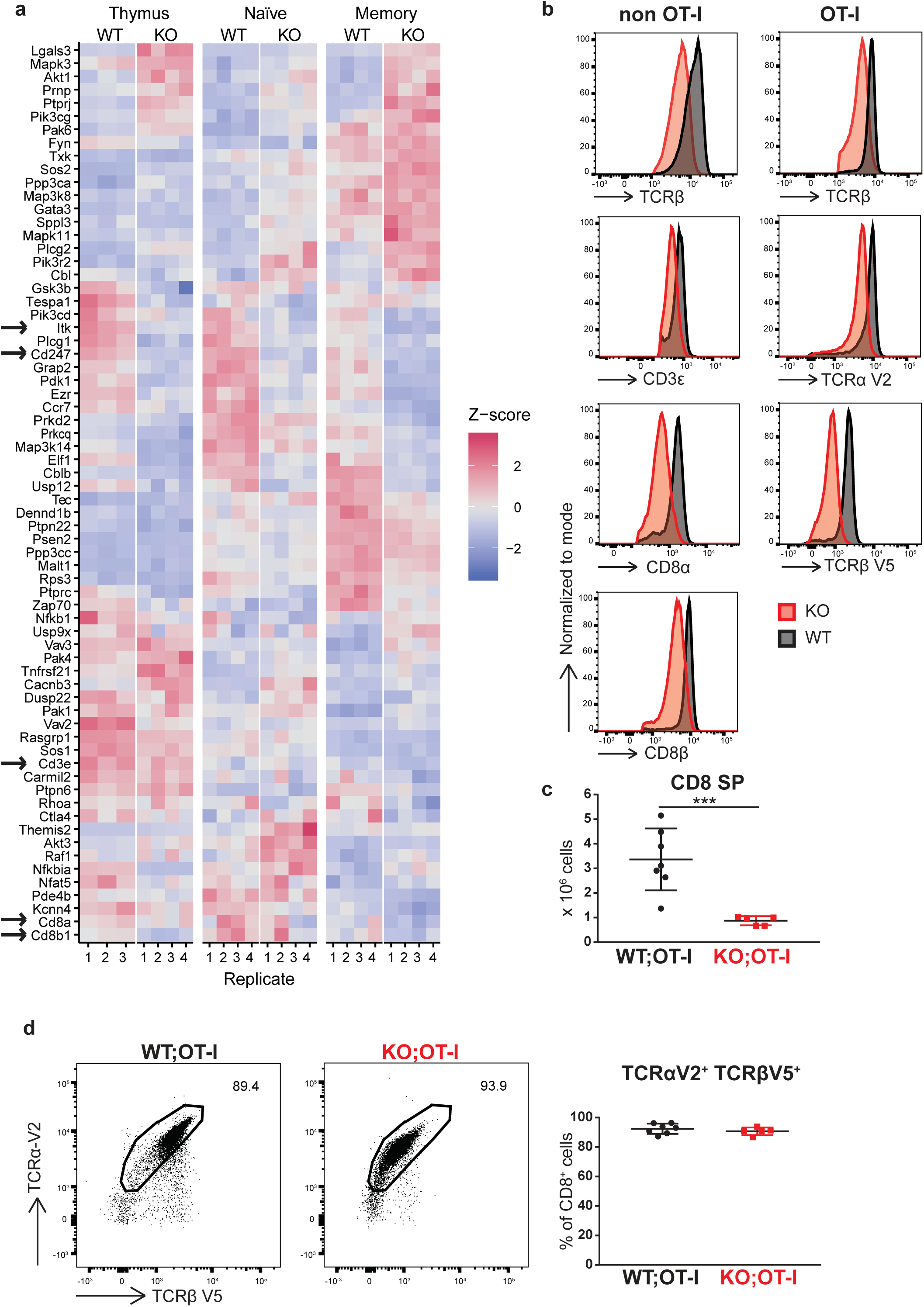
*Dot1L* ablation impairs TCR/CD3 expression. a) RNA expression of TCR signaling genes, defined by differential expression between WT and *Dot1L*-KO, in any of the sorted CD4^−^CD8^+^CD3^+^ thymocytes (thymus), CD44^−^CD62L^+^ (naïve) CD8^+^ T cells and CD44^+^CD62L^+^ (memory) CD8^+^ T cells, four mice per genotype except for WT thymus where there are three mice. Genes are clustered based on z-score. Arrows indicate genes involved in the TCR complex and Itk. Z-score is calculated row-wise within genes between samples. b) Expression of TCRβ, CD3e, TCRα V2, TCRβ V5, CD8α and CD8β on CD8^+^ T cells in spleen from *Lck-*Cre*;Dot1L* and *Lck-*Cre*;Dot1L;OT-I* mice. c) Absolute number of CD4^−^CD8^+^CD3^+^ thymocytes in WT and KO mice from *Lck-*Cre*;Dot1L;OT-I* background, data from two individual experiments with 3-4 mice per genotype, represented as mean±SD. d) Representative plot and quantification of TCRα V2 and TCRβ V5 expression on CD8^+^ T cells from spleen. Data from two experiments with 2-4 mice per genotype, represented as mean ±SD.

### *Dot1L*-KO T_AIM_ cells display memory features but are functionally impaired *in vitro*

A hallmark of memory T cells is that they proliferate faster and produce IFNγ rapidly upon stimulation compared to naïve T cells ^51^. Likewise, innate and virtual memory T cells rapidly produce IFNγ upon TCR stimulation ^38, 44^. To determine the proliferative potential of *Dot1L*-KO T_AIM_ cells, we activated B-cell depleted, CFSE-labeled splenocytes *in vitro* with anti-CD3 and anti-CD28 antibodies. Proliferation was determined by CFSE dilution. *Dot1L*-KO CD8^+^ T cells proliferated faster than WT cells (Fig. 4a). In addition, they rapidly became activated CD44^+^CD62L^−^ (Sup Fig. 4a). However, in contrast to WT, KO CD8^+^ T cells failed to produce IFNγ (Fig. 4b and Sup Fig. 4b), and expressed lower levels of CD69, a marker of activation (Sup Fig. 4c). This indicates a functional impairment of *Dot1L*-KO T cells. To test their intrinsic competence to produce IFNγ, *Dot1L*-KO T cells were stimulated in a TCR-independent manner with PMA and Ionomycin. Intracellular staining for IFNγ revealed that the percentage of stimulated CD8^+^ T cells producing IFNγ was 1.9-fold higher in KO as compared to WT (Sup Fig. 4d). This suggests that *Dot1L*-KO CD8^+^ T cells only partially respond to TCR stimulation, but that they have the intrinsic capacity to produce IFNγ. The responders in the WT population likely include the naturally existing virtual memory population (Fig. 1). Taken together, *Dot1L* KO T_AIM_ cells displayed some memory-associated features but simultaneously are functionally impaired.

**Figure 4:**
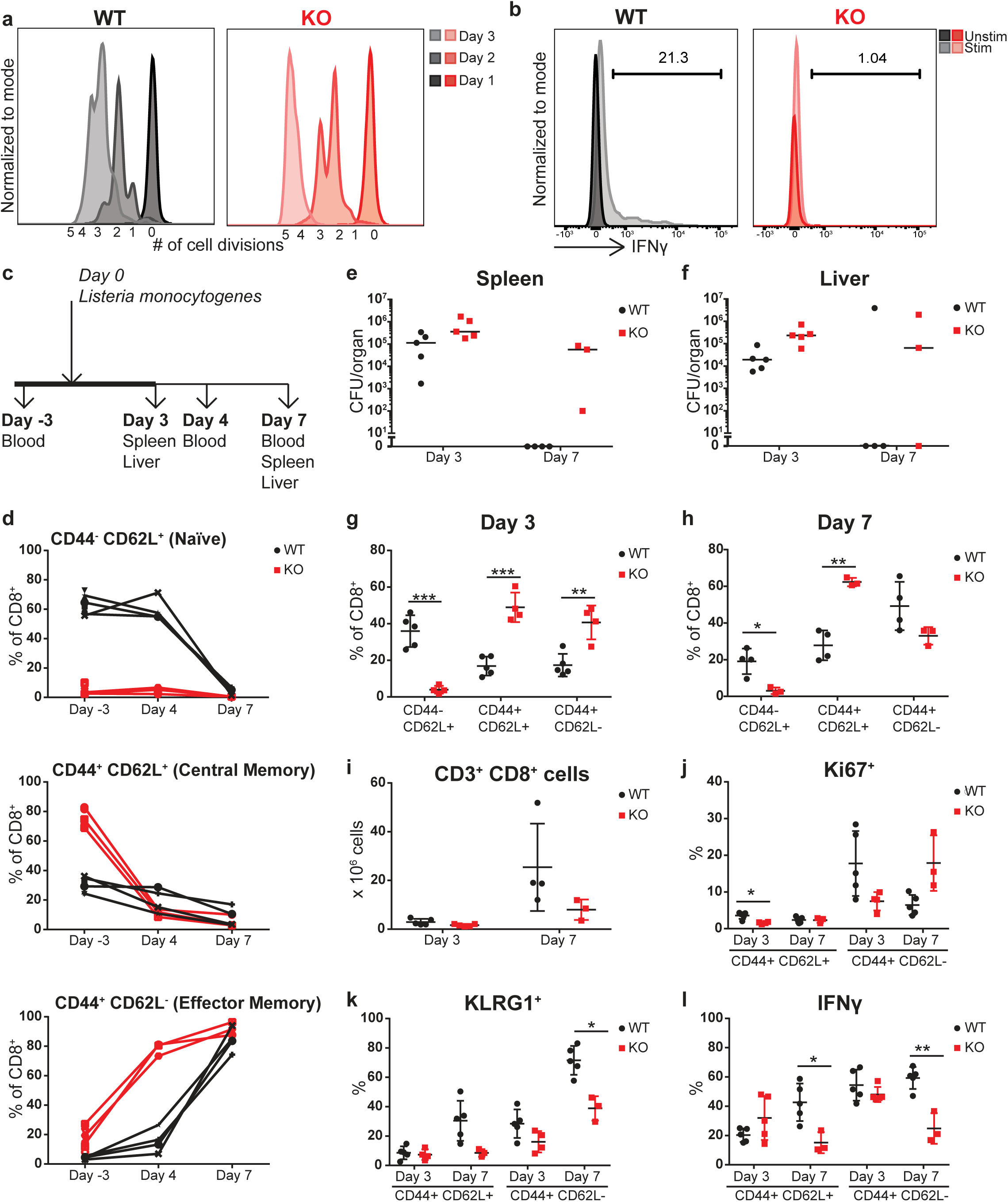
*Dot1L*-KO T_AIM_ cells have an impaired immune response in vitro and in vivo. a) Analysis of proliferation of CFSE-labeled WT and *Dot1L*-KO CD8^+^ T cells stimulated with α-CD3/ α-CD28 in vitro. b) IFNγ production by CD8^+^ T cells after stimulation with α-CD3/α-CD28 for 3 days and Golgi-plug incubation for four hours. c) Outline of immune challenge experiment. On day 0 mice were injected intravenously with a sub-lethal dose of *L. monocytogenes*. Blood was taken at day -3, day 4 and day 7 past injection. At day 3 and 7 past injection mice were sacrificed and spleen and liver were isolated for analysis. On day 3 five mice per genotype were sacrificed, on day 7 3-5 mice per genotype. d) Percentage of CD44 CD62L subsets of CD8^+^ T cells in blood. e-f) Clearance of *L. monocytogenes* in spleen and liver defined by the number of colony forming units (CFU) per organ. Bars indicate median. g) Percentage of CD44 CD62L subsets in CD8^+^ T cells from spleen at day 3 and h) day 7. i) absolute number of CD3^+^CD8^+^ T cells in spleen at day 3 and day 7. j) Intracellular Ki67 positive cells in CD8^+^ splenocytes. k) Percentage of KLRG1^+^ in CD8^+^ T cells from the spleen. l) Percentage of IFNγ-producing cells in CD8^+^ T cells from spleen after four hours stimulation with PMA/Ionomycin. g-l) Data is represented as mean ±SD.

### DOT1L is essential for mounting an effective immune response *in vivo*

To determine the *in vivo* immune responsiveness of *Dot1L-*KO T cells, mice were challenged with a sub-lethal dose of *Listeria monocytogenes*. The immune response was monitored in the blood at day -3, 4 and 7 post injection (Fig. 4c). Already at day 4 most CD8^+^ T cells in the blood had differentiated towards an effector-phenotype (CD44^+^CD62L^−^) in KO mice (Fig. 4d), which is in line with the *in vitro* stimulation data. However, in accordance with the peak of a normal immune response initiated from naïve T cells ^52^, the peak of activated effectors in WT mice was reached only after seven days. At day 3 and day 7 post injection, mice were sacrificed and spleen and liver were used to determine clearance of the *Listeria* by counting colony-forming units. Despite the fact that in KO at day 3 most T_AIM_ cells had differentiated into activated T cells, the *Listeria* infection was not cleared. At day 7 complete clearance was observed in the spleen from WT-mice, which is in accordance with literature ^52^; however, the *Dot1L* KO mice failed to clear *Listeria* (Fig. 4e-f). In order to investigate the reason for the compromised clearance in KO, despite the rapid T_EFF_ formation in the blood upon infection, we examined the splenic CD8^+^ T cells subsets in more detail. In WT mice the peak of activated (CD44^+^CD62L^−^) CD8^+^ T cells was at day 7, whereas in *Dot1L* KO mice this was at day 3 (Fig. 4g-h and Sup Fig. 4e). However, the absolute numbers of total CD8^+^ T cells in KO on day 3 were not higher than in WT. Furthermore, in WT the number of CD8^+^ T cells at day 7 had increased 8.6-fold, whereas in KO this was only 4.9-fold, suggesting a lack of expansion despite the acquisition of an effector phenotype (Fig. 4i). Indeed, staining for Ki67, a marker of proliferation showed that at day 3, the frequency of Ki67^+^ CD44^+^CD62L^−^CD8^+^ T cells was not increased in KO cells compared to WT (Fig. 4j). Besides the reduced proliferative capacity *in vivo*, even up to day 7, the KO CD44^+^CD62L^−^ CD8^+^ T cells showed impaired induction of the effector hallmarks KLRG1, IFNγ and CD49d (Fig. 4k-l, Sup Fig. 4f). Together these data indicate that upon *in vivo* challenge with *Listeria monocytogenes*, compromised clearance in KO mice relates to a reduced capacity to proliferate and terminally differentiate. In conclusion, DOT1L is essential in controlling the proliferation and effector function of T cells.

### DOT1L is required for maintenance of epigenetic integrity of CD8^+^ T cells

In order to mechanistically understand the functional impairment of *Dot1L*-KO CD8^+^ T_AIM_ cells *in vivo* and how this relates to the epigenome, we performed H3K79me2 ChIP-Seq on sorted CD8^+^ T cell populations and compared it to the RNA-Seq data. RNA-Seq analyses from CD44^−^CD62L^+^ (T_N_) and CD44^+^CD62L^+^ (T_CM_) cells revealed that genes that were upregulated in *Dot1L* KO were biased towards being lowly expressed, whereas downregulated genes tended to have higher expression level (Fig. 5a, left panel). To assess how the transcriptome changes in T cells and the transcriptional bias are related to the chromatin-modifying function of DOT1L, we compared the level of H3K79me2 at the 5’ end of genes in WT cells with the mRNA expression changes caused by the loss of DOT1L. H3K79me2 was enriched from the transcription start site into the first internal intron of transcribed genes (Sup Fig. 5a), as reported for human cells ^53^. Further analysis showed that most of the lowly-expressed genes upregulated in *Dot1L* KO contained no or very low H3K79me2 in WT cells (Fig. 5a right panel and Sup Fig. 5b). Therefore, these genes were not direct targets of DOT1L and are likely to be indirectly controlled by DOT1L. In contrast, most of the more highly-expressed genes downregulated in *Dot1L* KO were marked by H3K79me2 (Fig. 5a right panel and Sup Fig. 5b). This demonstrates that in normal CD8^+^ T cells DOT1L-mediated H3K79 methylation generally marks expressed genes, but only a subset of the methylated genes needs H3K79 methylation for maintaining full expression levels. Thus, in normal T cells DOT1L does not act as a transcriptional switch but rather seems to be required for transcription maintenance of a subset of methylated genes that are already on and it indirectly promotes repression of genes.

**Figure 5:**
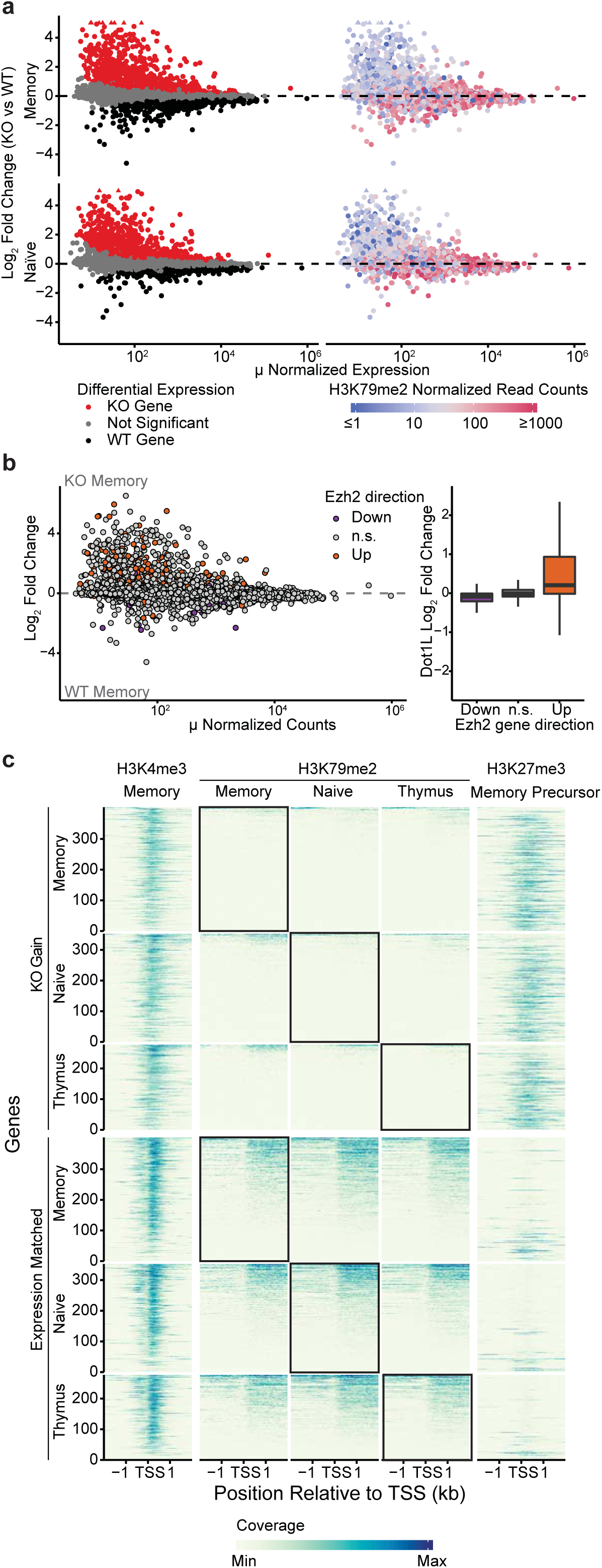
DOT1L is required for maintenance of the epigenetic identity of CD8^+^ T cells. a) MA plot indicating differentially expressed genes between *Dot1L* KO and WT in sorted CD44^−^CD62L^+^ (naïve) CD8^+^ T cells and CD44^+^CD62L^+^ (memory) CD8^+^ T cells (left). Genes significantly upregulated (red) and downregulated (black) in Dot1L-KO are indicated. Genes were considered differentially expressed when FDR <0.01. Genes outside the limit of the y-axis are indicated with triangles. Right panel shows H3K79me2 normalized read counts of these genes. Genes with low H3K79me2 read counts are blue, genes with high H3K79me2 are red. b) MA plot of gene expression between WT and Dot1L-KO memory CD8^+^ T cells, with genes differential between activated WT and activated *Ezh2*-KO CD8^+^ T cells indicated in orange (derepressed in *Ezh2* KO) and blue (downregulated in *Ezh2* KO). Box plot notes effect size in *Dot1L* KO vs WT RNA-Seq data based on the indicated differential gene expression category in *Ezh2* KO data. c) H3K4me3, H3K79me2 and H3K27me3 ChIP-Seq data around transcription start sites from genes that are upregulated in Dot1L KO (KO Gain) and genes that did not change expression (Expression Matched). Coverage was calculated as reads per genomic content, cutoff at the 0.995^th^ quantile and rescaled to a maximum of 1.

Amongst the differentially expressed genes marked with H3K79me2, we searched for candidate direct target genes that could mediate the role of DOT1L in terminal T cell differentiation. We were especially interested in factors that could explain the prominent gene derepression in *Dot1L*-KO cells. To this end we selected genes that were significantly downregulated in KO and had harbored H3K79me2 at the 5’ end of the gene, both in T_N_ and T_CM_. We further narrowed the list down to genes that were annotated as “negative regulator of transcription by RNA polymerase II”, resulting in 14 transcriptional regulators. Among those, *Ezh2* emerged as a potentially relevant target of DOT1L that could explain part of the derepression of genes in *Dot1L*-KO cells (Table 1). EZH2 is part of the Polycomb-repressive complex 2 (PRC2), which deposits H3K27me3 ^54–59^, a mark involved in repression of developmentally-regulated genes and in switching off naïve and memory genes during terminal differentiation of effector CD8^+^ T cells ^40, 60^. Although the change in *Ezh2* mRNA expression was modest between WT and KO (Sup Fig. 5c), the *Ezh2* gene was H3K79me2 methylated (Sup Fig. 5d), and *Dot1L*-KO mice share several T-cell phenotypes with *Ezh2*-KO mice ^40, 60^, although this is dependent on the mouse model used ^61–63^. To further investigate the idea that misregulation of PRC2 targets could be one of the downstream consequences of loss of DOT1L in CD8^+^ T cells, we compared the gene expression changes in an *Ezh2*-KO model ^61^ with those seen in *Dot1L*-KO CD8^+^ T cells. This revealed substantial overlap between the derepressed genes in the two models (Fig. 5b), suggesting a functional connection between two seemingly opposing epigenetic pathways. Furthermore, we determined H3K27me3 scores based on previous ChIP-Seq studies ^64^ and compared them with the gene expression in WT and *Dot1L*-KO SP CD8^+^ thymocytes and peripheral CD8^+^ T cells. This analysis showed that the genes that were upregulated in *Dot1L*-KO and lack H3K79me2 in WT were strongly enriched for H3K27me3 in WT memory precursor CD8^+^ T cells (Fig. 5c). As a control, expression matched non-differentially expressed genes were not enriched for H3K27me3 (Fig. 5c). Taken together, these findings suggest that one of the consequences of loss of DOT1L-mediated H3K79me2 is derepression of a subset of PRC2 targets that are normally actively repressed. This, together with the other transcriptional changes likely contributes to the perturbation of the epigenetic identity of CD8^+^ T_AIM_ cells, thereby compromising their ability to terminally differentiate *in vivo*.

**Table 1:** List of candidate negative regulators affected by DOT1L in T cells.

| Gene | Transcriptional regulator |
| --- | --- |
| *Hif1a* | yes |
| *Srebf2* | yes |
| *Lef1* | yes |
| *Notch1* | yes |
| *Mbd2* | yes |
| *Ezr* | no |
| *Zeb1* | yes |
| *Usp3* | yes |
| *Satb1* | yes |
| *Fnip1* | no |
| *Phb* | no |
| *Ezh2* | yes |
| *Zbtb20* | yes |
| *Rps14* | no |
| *Ybx1* | yes |
| *Smad7* | no |
| *Rpl10* | no |
| *Smad4* | no |
| *Gatad2a* | yes |
| *Ski* | no |
| *Etv3* | yes |
| *Foxk1* | yes |

## Discussion

The histone methyltransferase DOT1L has emerged as a druggable target in MLL-rearranged leukemia and additional roles in cancer have been suggested ^22–24, 65–68^. This in combination with the availability of highly-specific DOT1L inhibitors make DOT1L a potential target for cancer therapy. However, the role of DOT1L in normal lymphocyte physiology remained unknown. Here we show that DOT1L, plays a central role in ensuring normal T cell differentiation. To our knowledge, DOT1L is the first reported activating histone methyltransferase with a causal role in T cell differentiation.

T cell differentiation is intimately linked to epigenetic programming, but the mechanistic role of epigenetic marks in steering T cell differentiation ^40, 60, 61, 63, 69, 70^ and thymic selection ^71–75^ has only recently become more clear. Here we observed that ablation of *Dot1L* in the T cell lineage differentially affected CD4^+^ and CD8^+^ T cells. Following deletion of *Dot1L* by *Lck-*Cre during early thymocyte development, CD4^+^ T cells were strongly reduced, while the number of CD8^+^ T cells was increased. The fact that a substantial fraction of the remaining CD4^+^ T cells had not lost DOT1L activity indicates that the CD4^+^ compartment and especially regulatory T cells depend on DOT1L for their normal development. Further research exploring the dependency of CD4^+^ T cells on DOT1L may provide novel strategies for immune modulation and treatment of CD4^+^ T cell malignancies ^76, 77^.

In CD8^+^ T cells, loss of DOT1L resulted in a massive gain of CD44^+^CD62L^+^ memory-like cells associated with a simultaneous loss of the naïve compartment. These cells start to acquire memory features intrathymically, express a diverse un-skewed TCRβ repertoire, and lack expression of CD49d. Taken together, this suggests that these memory-like CD8^+^ T cells arose independently of foreign antigens, leading us to designate these cells as antigen-independent memory-like T_AIM_ CD8^+^ cells. Importantly, such unconventional memory-phenotype cells constitute a substantial (15-25%) fraction of the peripheral CD8^+^ T cell compartment in WT mice and humans ^4, 35, 44, 78, 79^. The fraction of unconventional memory-phenotype T cells further increases with age ^80, 81^. The biological role of antigen-inexperienced memory CD8^+^ T cells is still not completely understood, but these cells have been observed to respond more rapidly to TCR activation than T_N_ cells and they have been suggested to provide by-stander protection against infection in an antigen-independent way ^34, 44^. However, in older mice, virtual memory cells have been reported to lose their proliferative potential and acquire characteristics of senescence ^82^. The mechanism of unconventional memory-phenotype differentiation is also poorly understood ^2^. In some mouse models CD8^+^ T cells differentiate upon excess production of IL-4 by iNKT or γδ T cells ^3, 39, 83–86^. In contrast, other mouse models report that antigen inexperienced memory CD8^+^ T cell differentiation is cell intrinsic ^87, 88^. Here we show that DOT1L plays an important role in T_AIM_ CD8^+^ cell differentiation by cell-intrinsic mechanisms.

It has been established in several independent mouse models that the quality of TCR signaling closely relates to the formation of T_AIM_ cells ^34, 43, 44^. Our results show that loss of DOT1L leads to reduced surface expression of the CD3/TCR complex and co-receptors. This phenotype is likely related to the reduced expression of CD3ζ (Cd247), a target of DOT1L and a rate limiting molecule for assembly and transport of TCR/CD3 complexes to the cell surface ^1, 45–48^. The failure to upregulate the TCR/CD3 complex upon positive selection likely prohibits differentiation of conventional T_N_ and supports formation of T_AIM_ cells. In addition, *Dot1L* ablation perturbed expression of TCR signaling genes, including *Itk* (IL2-inducible T cell kinase; a member of the Tec kinase family). Disruption of ITK signaling has also been reported to lead to antigen-independent T-cell differentiation ^49^. Together, this suggests that one of the key functions of DOT1L is to ensure adequate TCR surface expression and signaling to maintain naivety and prevent T_AIM_ cell differentiation. The discovery of DOT1L as a key player in preventing premature antigen-independent differentiation towards memory-type cells warrants further investigation and will aid in further uncovering the origin and regulation of this emerging and intriguing subset of the immune system.

Functionally, *Dot1L*-KO CD8^+^ T_AIM_ cells partially displayed features of memory cells *in vitro* but *in vivo* showed an impaired immune response against *Listeria monocytogenes*. The rapid differentiation towards effector cells suggests that *Dot1L*-KO T_AIM_ cells might execute a partial by-stander response, but DOT1L appears to be required to accomplish full effector function. Interestingly, similar features have been observed for virtual memory cells ^44^. The reduced TCR density and affected TCR signaling network in *Dot1L*-KO T cells likely further contribute to the impaired immune response.

How does DOT1L affect T cell differentiation at the chromatin level? Inspection of the transcriptome and epigenome provided evidence that DOT1L methylates transcriptionally active genes in T cells and positively affects gene expression. However, only a subset of the targets required DOT1L for maintenance of normal expression levels, which is in agreement with previous observations ^89, 90^. Why some genes depend on DOT1L/H3K79 methylation and others not is not known yet although a recent study indicates that in MLL-rearranged leukemic cell lines some genes harbor a 3’ enhancer located in the H3K79me2/3 marked genic region, which can make them more sensitive to loss of DOT1L ^89^. One of the genes that was H3K79 methylated and dependent on DOT1L in normal peripheral CD8^+^ T cells was *Ezh2* and EZH2/PRC2 targets were derepressed in the absence of DOT1L. Importantly, analysis of data from Kagoya et al. ^26^ showed that *Ezh2* expression is also reduced in human T cells in which DOT1L was inactivated not by deletion but by treatment with a DOT1L inhibitor (Sup Fig. 5e). This suggests that the epigenetic crosstalk that we uncovered in mice is evolutionarily conserved. Interestingly, recruitment of DOT1L to nucleosomes by its interaction partner AF10 has been shown to be negatively affected by methylation of H3K27 ^91^. Therefore, the crosstalk between DOT1L and PRC2 might involve mutual interactions. Restricting DOT1L to nucleosomes unmethylated at H3K27 might be one of the mechanisms by which H3K79me and H3K27me3 are directed to non-overlapping sites, as we also observed in T cells (Fig. 5C).

Derepression of some of the targets of PRC2 is just one of the consequences of loss of DOT1L. Besides *Ezh2*, DOT1L affects the expression of other genes, including several additional candidate transcription regulators (Table 1). In the future, it will be important to determine other mechanisms by which DOT1L affects the CD8^+^ T cell transcriptome to fully understand its central role in CD8^+^ T cell biology. Finally, we cannot exclude that DOT1L has additional methylation targets besides H3K79 that contribute to role of DOT1L in safeguarding T cell differentiation and effector functions. Although understanding the mechanisms in more detail will require further studies, the role of DOT1L in preventing premature differentiation and safeguarding the epigenetic identity is conserved in other lymphocyte subsets. In an accompanying study (Aslam et al), we observed in an independent mouse model that loss of *Dot1L* in B cells also led to premature differentiation, perturbed repression of PRC2 targets, and a compromised humoral immune response. Therefore, DOT1L is emerging as a central epigenetic regulator of lymphocyte differentiation and functionality.

In conclusion, we identify H3K79 methylation by DOT1L as an activating epigenetic mark critical for CD8^+^ T cell differentiation and maintenance of epigenetic identity. Further investigation into the central role of the druggable epigenetic writer DOT1L in lymphocytes is likely to provide novel strategies for immune modulations and disease intervention ^92–95^.

## Methods

### Mice

*Lck-*Cre*;Dot1L^fl/fl^* mice have been described elsewhere previously ^25^ and were based on the Dot1Ltm1a(KOMP)Wtsi line generated by the Wellcome Trust Sanger Institute (WTSI) and obtained from the KOMP Repository (www.komp.org) ^96^. Mice from this newly created *Lck-*Cre*;Dot1L* strain were breeding in a mendelian ratio and had no welfare issues. *Lck-*Cre*^+/−^;Dot1L^fl/fl^* (KO) mice were compared to *Lck-*Cre*^+/−^;Dot1L^wt/wt^* (WT) mice and were indicated with *Lck-*Cre*^+/−^;Dot1L^fl/wt^* (Het) mice in order to eliminate any Cre-specific effects ^97^. The *Lck-*Cre*;Dot1L^fl/fl^* mice were crossed with OT(B6J) mice (a kind gift from the Ton Schumacher group, originally from Jackson labs) to generate *Lck-*Cre*;Dot1L^fl/fl^;OT-I* mice. Both strains were bred in-house. Mice used for experiments were between 6 weeks and 8 months old and of both genders. For each individual experiment mice were matched for age and gender. Mice were housed under specific pathogen free (SPF) conditions at the animal laboratory facility of the Netherlands Cancer Institute (NKI; Amsterdam, Netherlands). All experiments were approved by the Animal Ethics Committee of the NKI and performed in accordance with institutional, national and European guidelines for animal care and use.

### Flow cytometry

Single cell suspensions were made from spleen and thymus. Erylysis was performed on blood and spleen samples. Cells were stained with fluorescently labeled antibodies in a 1:200 dilution unless otherwise indicated (Table 2). Of note, for OT-I tetramer stains the CD8 antibody clone 53-6.7 was used ^98^. For intracellular staining, cells were fixed and permeabilized using the Transcription Factor Buffer kit (Benton Dickinson). Antibodies for intracellular staining were diluted 1:200 in Perm/Wash buffer (Table 3). For H3K79me2 staining, cells were first stained with surface markers and fixed and permeabilized as described before. After fixation and permeabilization cells were washed with Perm/Wash containing 0.25% SDS. α-H3K79me2 (Millipore) was diluted 1:200 into Perm/Wash + 0.25% SDS and cells were incubated for 30 min. Cells were washed with Perm/Wash and incubated with the secondary antibodies Donkey anti-Rabbit AF555 (Thermo Scientific) or Goat-anti-Rabbit AF488 (Invitrogen) 1:1000 in Perm/Wash. Flow cytometry was performed using the LSR Fortessa (BD Biosciences) and data were analyzed with FlowJo software (Tree Star inc.) Histograms were smoothed.

**Table 2:** Antibodies for surface stains.

| **Antibody** | **Fluorescent label** | **Clone** | **Company** |
| --- | --- | --- | --- |
| CD3e | APC | 145-2C11 | BD |
| CD3e | AF488 | 145-2C11 | eBioscience |
| TCRβ | PB | H57-597 | BioLegend |
| TCRβ | PE | H57-597 | BD |
| CD4 | APC | RM4-5 | BioLegend |
| CD4 | BV785 | GK1.5 | BioLegend |
| CD8a | BV510 | 53-6.7 | BioLegend |
| CD8b | PerCpCy5.5 | YTS1567.7.7 | BioLegend |
| CD19 | APC-H7 | 1D3 | BD |
| CD19 | PE | 1D3 | BD |
| CD25 | APC | PC61 | BioLegend |
| CD49d | AF488 | R-12 | BioLegend |
| CD44 | APC-Cy7 | IM7 | BioLegend |
| CD44 | BV650 | IM7 | BioLegend |
| CD45 | PerCpCy5.5 | 30-F11 | BioLegend |
| CD62L | PE | MEL-14 | BD |
| CD62L | PE-Cy7 | MEL-14 | BioLegend |
| CD62L | FITC | MEL-14 | BD Pharmingen/eBioscience |
| CD69 | PE | H1.2F3 | BD |
| CD90.2 | FITC | 53-2.1 | BD |
| KLRG1 | BV421 | 2F1/KLRG1 | BioLegend |
| $\gamma\delta$TCR | BV605 | GL3 | BioLegend |
| TCRαV2 | PE | B20.1 | BD |
| TCRβV5 | APC | MR9-4 | eBioscience |
| OT-I tetramer | PE | NA | Schumacher group |
| CD1d tetramer (1:100) | AF488 | NA | NIH Tetramer Core Facility |
| LIVE/DEAD Fixable<br>Near-IR Dead Cell Stain<br>Kit | NA | NA | Thermo Fisher |
| Zombie NIR | NA | NA | BioLegend |
| Zombie Violet | NA | NA | BioLegend |

**Table 3:** Antibodies for intracellular stains.

| Antibody | Fluorescent label | Clone | Company |
| --- | --- | --- | --- |
| IFNγ | PE | XMG1.2 | BD |
| T-bet | PE-Cy7 | 4B10 | eBioscience |
| EOMES | AF488 | Dan11mag | eBioscience |
| EOMES | eF450 | Dan11mag | eBioscience |
| IFNγ | PE-Cy7 | XMG1.2 | eBioscience |
| Ki67 | PE | SolA15 | eBioscience |

### Immunohistochemistry

Thymus tissues were fixed in EAF (ethanol, acetic acid, formol saline) for 24 hours and subsequently embedded in paraffin. Immunohistochemistry was performed with H3K79me2 antibody ^99^as described in ^25^.

### PCR

Sorted CD4^+^ and CD8^+^ T cells were lysed in DirectPCR Lysis Reagent (mouse tail) (Viagen Biotech) with 1mg/ml proteinase K (Sigma). *Dot1L^fl/fl^* and *Dot1L*^Δ/Δ^ was detected by PCR using the following primers. Dot1L_FWD: GCAAGCCTACAGCCTTCATC, Dot1L_REV: CACCGGATAGTCTCAATAATCTCA and Dot1L_Δ: GAACCACAGGATGCTTCAG and MyTaq Red Mix (GC Biotech).

### *In vitro* stimulation

To determine cytokine production upon *in vitro* stimulation, splenocytes were stimulated with 20 ng/ml PMA (Sigma), 0.5 μg/ml Ionomcyin (Sigma) and 1μl/ml Golgi Plug protein transport inhibitor (Benton Dickinson) for 4 hours. Cells were stained as described above. To determine proliferation and IFNγ production upon *in vitro* TCR-mediated stimulation splenocytes were enriched for T cells using CD19 microbeads depletion (Miltenyi Biotec) on LS columns (Miltenyi Biotec). Cells were labeled with 0.5mM CFSE (Thermo Scientific) in order to measure proliferation and left unlabeled to determine IFNγ production and plated in a 96-wells plate coated with anti-CD3 (145-2C11) (BD) in complete medium (IMDM supplemented with 8% FCS, 100 mM pen/strep and 100 mM β-mercaptoethanol). Anti-CD28 (37.51) (BD) was added to the medium and cells were incubated at 37 °C with 5% CO_2_. At day 1 after CFSE labeling, cells were analyzed by flow cytometry to confirm the uptake of CFSE. In order to determine IFNγ production cells were incubated with Golgi Plug protein transport inhibitor (Benton Dickinson) for 4 hours.

### *In vivo* immunization with *Listeria monocytogenes*

*Listeria monocytogenes* strain LM-OVA was a gift from Ton Schumacher (NKI). Bacteria were grown overnight in Bacto Brain Heart Infusion, Porcine (BHI) medium (BD) to an absorbance at 600 nm of OD 0.9. A sub-lethal dose (10.000 colony forming units; CFUs) of *Listeria monocytogenes* in HBSS was injected intravenously into the mice. The number of bacteria injected was confirmed by growth on BHI agar plates (BD) with 25 μg/ml chloramphenicol. The immune response was followed in the blood by taking blood samples via tail vein puncture at day -3 and +4. The CFUs were determined for spleen and liver. Single-cell suspensions were made in PBS and the cells were lysed in 0.1 % IGEPAL. Homogenized suspensions were plated on BHI agar plates in three dilutions (undiluted, 1:10 and 1:100). After 48 hours colony counts were determined.

### Sort for RNA-Seq, ChIP-Seq and TCR-Seq

For cell sorting, thymus and spleen samples were homogenized. Erylysis was performed on spleen samples. Cells were stained with fluorescently labeled antibodies (1:200) (Table 4) and sorted on FACSAria Ilu (BD Biosciences), FACSAria Fusion (BD Biosciences) or MoFlo Astrios (Beckman Coulter). Cells were collected in tubes coated with FCS.

**Table 4:** Antibodies for cell sorting.

| Antibody | Fluorescent label | Clone | Company |
| --- | --- | --- | --- |
| CD3 | AF488 | 145-2C11 | eBioscience |
| CD4 | APC | RM4-5 | BioLegend |
| CD8 | PerCp-Cy5.5 | YTS1567.7.7 | BioLegend |
| CD44 | APC-Cy7 | IM7 | BioLegend |
| CD62L | PE | MEL-14 | BD biosciences |
| TCRβ | PE | H57-597 | BD |

### RNA-Seq sample preparation

Flow cytometrically purified cells were resuspended in Trizol (Ambion Life Technologies) and total RNA was extracted according to the manufacturer’s protocol. Quality and quantity of the total RNA was assessed by the 2100 Bioanalyzer using a Nano chip (Agilent). Only RNA samples having an RNA Integrity Number (RIN) > 8 were subjected to library generation.

### RNA-Seq library preparation

Strand-specific cDNA libraries were generated using the TruSeq Stranded mRNA sample preparation kit (Illumina) according to the manufacturer’s protocol. The libraries were analyzed for size and quantity of cDNAs on a 2100 Bioanalyzer using a DNA 7500 chip (Agilent), diluted and pooled in multiplex sequencing pools. The libraries were sequenced as 65 base single reads on a HiSeq2500 (Illumina).

### RNA-Seq data preprocessing

RNA-Seq reads were mapped to mm10 (Ensembl GRCm38) using TopHat with the arguments ‘--prefilter-multihits –no-coverage-search –bowtie1 –library-type fr-firststrand’ using a transcriptome index. Counts per gene were obtained using htseq-count with the options ‘-m union -s nò and Ensembl GRCm38.90 gene models. Analysis was restricted to genes that have least 20 counts in at least 4 samples, and at least 5 counts in 4 samples in specific contrasts, to exclude very low abundance genes. Differential expression analysis was performed on only relevant samples using DESeq2 and default arguments with the design set to either *Dot1L*KO status or cell type. Adaptive effect size shrinkage was performed with the ashr R package to control for the lower information content in low abundance transcripts. Genes were considered to be differentially expressed when the p-value of the negative binomial Wald test was below 0.01 after the Benjamini-Hochberg multiple testing correction. Sets of differentially expressed genes in indicated conditions were called ‘gene signatures’. An exception was made for the Ezh2 RNA-Seq data from He et al., ^61^ wherein the dispersion was estimated with a local fit, using ‘estimateDispersion’ function in DESeq2 with the argument ‘fitType = “local”’ and using an adjusted p-value cutoff of 0.05, to increase the number of detected differential genes. Principal component analysis was performed using the ‘prcomp’ function on variance stabilizing transformed data of all samples with the ‘vst’ function from the DESeq2 package on all samples using default arguments. For analyses where we performed expression matching, we chose genes with an absolute log2 fold changes less than 0.1 and false discovery rate corrected p-values above 0.05 that were closest in mean expression to each of the genes being matched without replacement.

### ChIP-Seq sample preparation

Sorted cells were centrifuged at 500 rcf. The pellet was resuspended in IMDM containing 2% FCS and formaldehyde (Sigma) was added to a final concentration of 1%. After 10 min incubation at RT glycine (final concentration 125 mM) was added and incubated for 5 min. Cells were washed twice with ice-cold PBS containing Complete, EDTA free, protein inhibitor cocktail (PIC) (Roche). Cross-linked cell pellets were stored at −80 °C. Pellets were resuspended in cold Nuclei lysis buffer (50mM Tris-HCl pH 8.0, 10mM EDTA pH8.0, 1%SDS) + PIC and incubated for at least 10 min. Cells were sonicated with PICO to an average length of 200-500bp using 30s on/ 30s off for 3 min. After centrifugation at high speed debris was removed and 9x volume of ChIP dilution buffer (50mM Tris-HCl pH8, 0.167M NaCl, 1.1% Triton X-100, 0.11% sodium deoxycholate) + PIC and 5x volume of RIPA-150(50mM Tris-HCl pH8, 0.15M NaCl, 1mM EDTA pH8, 0.1% SDS, 1% Triton X-100, 0.1% sodium deoxycholate) + PIC was added. Shearing efficiency was confirmed by reverse crosslinking the chromatin and checking the size on agarose gel. Chromatin was pre-cleared by adding ProteinG Dynabeads (Life Technologies) and rotation for 1 hour at 4 °C. After the beads were removed 2μl H3K79me1, 2μl H3K79me2 (NL59, Merck Millipore) and 1μl H3K4me3 (ab8580, Abcam) were added and incubated overnight at 4°C. ProteinG Dynabeads were added to the IP and incubated for 3 hours at 4 °C. Beads with bound immune complexes were subsequently washed with RIPA-150, 2 times RIPA-500 (50mM Tris-HCl pH8, 0.5M NaCl, 1mM EDTA pH8, 0.1% SDS, 1% Triton X-100, 0.1% sodium deoxycholate), 2 times RIPA-LiCl (50mM Tris-HCl pH8, 1mM EDTA pH8, 1% Nonidet P-40, 0.7% sodium deoxycholate, 0.5M LiCl2) and TE. Beads were resuspended in 150 μl Direct elution buffer (10mM Tris-HCl pH8, 0.3M NaCl, 5mM EDTA pH8, 0.5%SDS) and incubated overnight at 65 °C and input samples were included. Supernatant was transferred to a new tub and 1μl RNAse A (Sigma) and 3 μl ProtK (Sigma) was added per sample and incubated at 55 °C for 1 hour. DNA was purified using Qiagen purification columns.

### ChIP-Seq Library preparation

Library preparation was done using KAPA LTP Library preparation kit using the manufacturer’s protocol with slight modifications. Briefly, after end-repair and A-tailing adaptor were ligated followed by Solid Phase Reverisble Immobilization (SPRI) clean-up. Libraries were amplified by PCR and fragments between 250-450 bp were selected using AMPure XP beads (Beckman Coultier). The libraries were analyzed for size and quantity of DNAs on a 2100 Bioanalyzer using a High Sensitivity DNA Kit (Agilent), diluted and pooled in multiplex sequencing pools. The libraries were sequenced as 65 base single reads on a HiSeq2500 (Illumina).

### ChIP-Seq data preprocessing

ChIP-Seq samples were mapped to mm10 (Ensembl GRCm38) using BWA-MEM with the option ‘-M’ and filtered for reads with a mapping quality higher than 37. Duplicate reads were removed using MarkDuplicates from the Picard toolset with ‘VALIDATION_STRINGENCY=LENIENT’ and ‘REMOVE_DUPLICATES=false’ as argument. Low quality reads with any FLAG bit set to 1804 were removed. Bigwig tracks were generated by using bamCoverage from deepTools using the following arguments: ‘-of bigwig –binsize 25 –normalizeUsing RPGC –ignoreForNormalization chrM – effectiveGenomeSize 2652783500’. For visualization of heatmaps and genomic tracks, bigwig files were loaded into R using the ‘import.bw()’ function from the rtracklayer R package. TSSs for heatmaps were taken from Ensembl GRCm38.90 gene models by taking the first base pair of the 5’ UTR of transcripts. When such annotation was missing, the most 5’ position of the first exon was taken.

### Defining DOT1L targets involved in transcription regulation

Genes were selected that are down in KO with adjusted p-value < 0.01, with H3K79me2 normalized reads > 20 and the GO annotation “negative regulator of transcription by RNA polymerase II”.

### TCRβ-Seq

Between 100,000 and 1,100,000 sorted cells were used for TCRβ-Seq. DNA was extracted using Qiagen DNA isolation kit (Qiagen) and collected in 50 μl TE buffer. The concentration was determined with Nanodrop. Samples were sequenced and processed by Adaptive Biotechnologies using immunoSEQ mm TCRB Service (Survey).

### Statistics

Statistical analyses were performed using Prism 7 (Graphpad). Data are presented as means ±SD unless otherwise indicated in the figure legends. The unpaired Student’s t-test with two-tailed distributions was used for statistical analyses. A p-value <0.05 was considered statistically significant. * p < 0.05, ** p < 0.01, *** p < 0.001.

## Acknowledgements

We thank the NKI animal pathology facility for histology and immunohistochemistry, as well as advice, the NKI genomics core facility for library preparations and sequencing, the NKI Flow Cytometry facility for assistance, the NKI Animal Laboratory Intervention Unit for performing immunization experiments, and the caretakers of the NKI Animal Laboratory facility for assistance and excellent animal care. The following reagent(s) was/were obtained through the NIH Tetramer Core Facility: CD1d-PBS57 and CD1d-unloaded tetramer. We thank Paul van den Berk for advice and Cheyenne Seerden for help with the flow cytometry experiments, Ton Schumacher and group for OT-I mice and OT-I tetramer, and Jan-Hermen Dannenberg for providing the Lck-Cre mouse model. We thank Muddassir Malik for critically reading the manuscript.

This work was supported by the Dutch Cancer Society (NKI2014-7232 to FvL and HJ) and the Netherlands Organisation for Scientific Research (NWO-VICI-016.130.627 to FvL; ZonMW Top 91213018 to HJ; ZonMW Top91218022 to FvL and HJ). The funders had no role in study design, data collection and interpretation, or the decision to submit the work for publication.

## Author contributions

EMK-M, MAA, MFA, FvL and HJ jointly designed the experiments

EMK-M, MAA, MFA jointly carried out the experiments

CMM, HV, CL, and SH generated the mouse model and performed the initial analyses on thymocytes

TvW performed genotyping and ChIP experiments

TvdB, MAA, TK, MFA, and EdW performed bioinformatics analyses

DvD and JMMdH performed the initial characterization of splenocytes

TA and JB contributed to the immunization experiments

FvL and HJ designed and supervised the studies.

EMK-M, MAA, MFA, FvL and HJ wrote the manuscript

